# Modelling the impact of rainfall events on increasing *Vibrio vulnificus* concentrations in Pacific oysters (*Crassostrea gigas*)

**DOI:** 10.1101/2023.08.29.555425

**Authors:** Graham C. Fletcher, Roland Taylor, Duncan Hedderley

## Abstract

*Vibrio vulnificus* occurs naturally in seawater and causes debilitating, often fatal illnesses, particularly in people with underlying health issues such as liver disease. The illness can occur when raw molluscan shellfish that have bio-accumulated the organism are consumed. New Zealand seafood is not known to have caused any illnesses, although there have been wound infections. This study sought to better understand the effect of environmental conditions on concentrations of *V. vulnificus* in Pacific oysters in three harbours over the summer months of 2016–19. Fortnightly sampling at two harbours only once gave a most probable number (MPN) of >10 per g of oyster meat, while the third harbour regularly produced much higher counts (up to 220,000 per g). From 2017–19, weekly samples from three sites in this harbour (four in 2018) were tested. Eleven peaks in concentration were observed, all when seawater temperatures exceeded 20°C and after heavy rainfall had reduced the seawater salinity, usually to <25‰ from the average of 32‰. A fitted structural equation model with temperature and salinity terms accounted for 60% of the variance in concentrations and rates of decline after concentration peaks averaged 1.05 log_10_ MPN per week.

The rate of decline was highly variable so microbiological testing would be required to confirm this rate for use in food safety management. However, the results of the study will enable better risk management.

**Importance:** *Vibrio vulnificus* is a deadly bacterium that naturally occurs in some seawater and can be concentrated in shellfish such as oysters. Consumers with underlying health issues who eat contaminated shellfish may suffer serious illnesses. New Zealand shellfish have never been known to cause such illnesses although the species is present in our seawater. The strains present may not be able to cause foodborne illness. We found that *V. vulnificus* could be present in high concentrations in farmed Pacific oysters in one of three studied harbours. This happened when seawater temperatures were warm and heavy rainfall reduced salinity. We developed a model based on seawater temperature and salinity that would be able to predict concentrations in shellfish, at least in this harbour. This will help industry and regulators manage the food safety risk should this organism become a public health issue.

**Tweet:** We modelled concentrations of *Vibrio vulnificus* in Pacific oysters after flood events and showed that seawater salinity and temperatures in previous 168 h affected them.

## Introduction

*Vibrio vulnificus* is an estuarine halophilic bacterium that can cause serious human diseases. There are increasing numbers of reported cases and a relatively high death rate (1, 2). The organism can be associated with gastroenteritis, but the two more serious diseases are primary septicaemia (foodborne) and necrotising wound infections (3, 4, 5). Serious disease is relatively rare, with the organism being an opportunistic pathogen rather than having human hosts as a primary vehicle of transmission (6). Patients almost invariably have pre-existing health issues and liver disease is a strong predictor of infection and death (7, 8). A United States review (1988–96) showed 48% of cases were foodborne with consumption of raw oysters being implicated in 96% of the septicaemia cases. Consumption of clams and shrimp were also implicated (7) and other outbreaks have been associated with eating crab (9). Risk of illness appears to be increasing with climate change (10).

Studies modelling climate change suggest that with ocean warming, *V. vulnificus* densities in seawater are likely to increase by up to 40% by 2100. However, climate change is also likely to increase precipitation in some areas with more incidences of extreme daily rainfalls (11) that will reduce estuarine salinities.

Like most *Vibrio* organisms, *V. vulnificus* is a natural part of the marine ecosystem and its presence is not due to contamination from terrestrial sources. However, concentrations are affected by environmental conditions, particularly seawater temperature and salinity. Numbers increase when temperatures exceed 18°C (5) and the organism enters a viable but non-culturable state at temperatures below 10°C (12). They are reported to occur in salinities between 0.2 and 30‰ with optimum salinities reported to be between 10 and 18‰ (4). Motes *et al*. (14) reported that salinities above 25‰ had a negative effect on *V. vulnificus* numbers, and on the Atlantic Coast (73% of oyster samples exceeding 26‰) the only sample with high numbers was collected in North Carolina after a late summer flood had reduced the salinity. However, the organism has often been found in New Zealand shellfish grown in marine and estuarine environments where the salinity is typically above 30‰ (14). The changing climate is resulting in more floods in New Zealand (15, 16), which will mean more periods of low seawater salinity (14). A recent review stated that “the paucity of high quality data from geographically diverse regions (other than the United States of America) probably represents the most pressing limitation for risk assessment efforts” for *Vibrio parahaemolyticus* and *V. vulnificus* (17)

New Zealand has had wound infections but has never had a confirmed case of foodborne *V. vulnificus* infection (18). It is a serious disease with the highest case fatality rate (∼50%) of any foodborne pathogen (6), so it is unlikely that many cases would be missed. In a 3-year study (2009– 12) of the organism’s prevalence in commercially farmed bivalve shellfish (13), *V. vulnificus* was found in 14% of Pacific oyster (*Crassostrea gigas*) samples (n = 235) but not in any Greenshell™ mussel (*Perna canaliculus*) or dredge oyster (*Ostrea chilenses*) samples (n= 55 and 21, respectively). Elevated concentrations (most probable numbers (MPN) exceeding 30 g^−1^) were only recorded during one (2011) of the three summers covered in the study when high numbers (up to 9300 MPN g^−1^) were recorded at four geographically distinct sites. Subsequent sampling of Pacific oysters each year for various reasons did not detect elevated concentrations until high numbers were detected in 2016 (Figure 1 derived from data presented by King *et al*. (19)). Analysis of *V. vulnificus* suggested that as well as being positively correlated to surface seawater temperature (SST), *V. vulnificus* concentrations were weakly negatively correlated with salinity, with lower salinities giving increasing concentrations (19). Further, in Harbour A of the study of Cruz *et al*. (13), *V. vulnificus* numbers were increasingly negatively correlated with salinity over the previous 10 days prior to sampling and increasingly positively correlated with rainfall and SST recorded in the previous 8 or 10 days before sampling, respectively. The current study was carried out to further evaluate and attempt to model this relationship.

**Figure 1.**
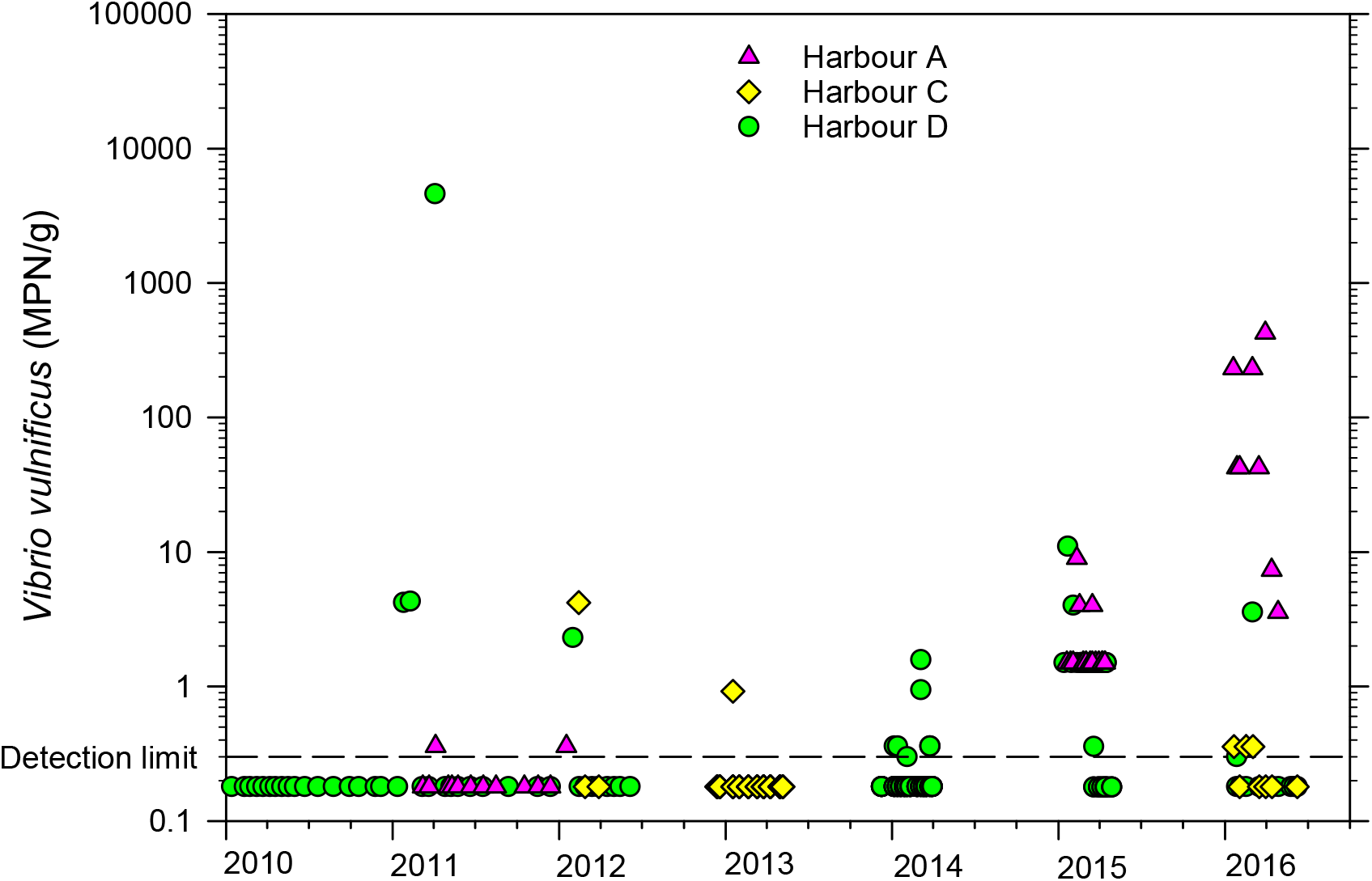
*Vibrio vulnificus* concentrations detected in Pacific oysters collected from three New Zealand growing harbours between 2010 and 2016.

## Materials and Methods

### Sampling programme

In 2016, a fortnightly monitoring programme included samples of Pacific oysters collected from three harbours (A, C and D) between 19 January and 7 June (20). As results showed high numbers of *V. vulnificus* in Harbour A, three Pacific oyster–growing sites (Figure 2) were selected on a commercial farm in the upper reaches of this harbour, designated Sites 7, 11 and 12. Weekly samples were collected between January and June for the next 3 years (2017–19). In 2018, samples were also collected at another farm closer to the mouth of Harbour A (Site 6 in Figure 2).

**Figure 2.**
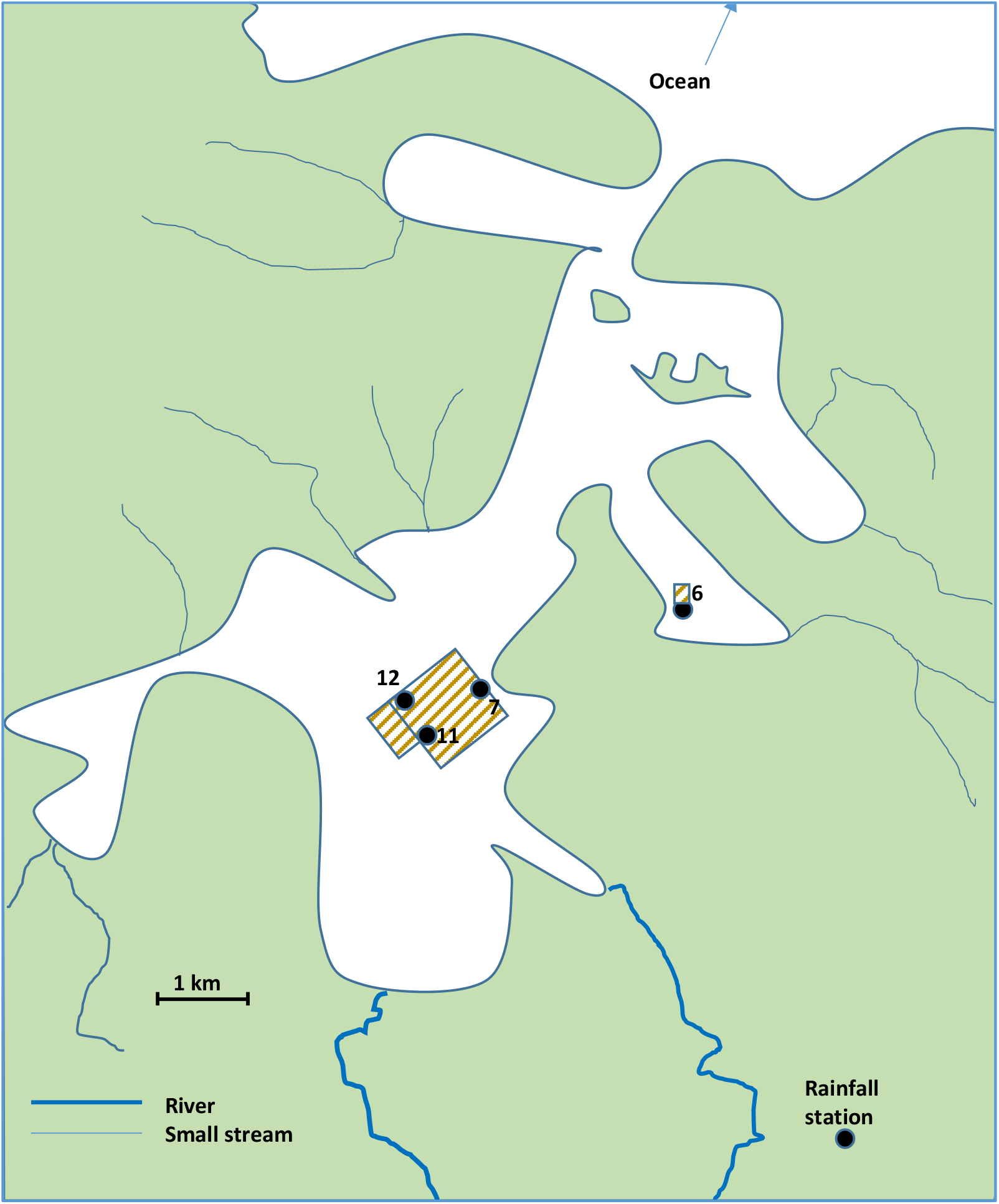
Schematic diagram of Harbour A showing sampling sites on two oyster farms (stripes) and the rainfall station. Temperatures and salinities were continuously recorded at Site 11, while all sampling sites had spot measurements when samples were collected.

Fortnightly samples were also collected from Harbours C and D for baseline comparisons. At each site, SST was measured with a thermometer when collecting the samples and a seawater sample was collected for subsequent salinity testing using a refractometer (Atago S/Mill, Japan). Additionally, the oyster farmers deployed a monitoring buoy at Site 11 recording SST and salinity at 15 min intervals and the buoy owner (Water and Atmosphere Information Limited, www.wai.co.nz) provided access to the data for the duration of the study. Rainfall data were obtained from the NIWA website (https://cliflo.niwa.co.New Zealand/) at the closest monitored sites.

Samples were placed in a polystyrene box containing but not in contact with ice and transported to the laboratory. Samples from Harbour A (the most distant site from the laboratory) took a mean of 24 h to arrive (maximum = 33 h with an exception on 10 May 2016) and arriving with a mean temperature of 12.4°C (range = 4.5–21°C). Although these times and temperatures (e. g. in the worst case a sample arriving at 20°C with a 33-h delivery time on 15 February 2017) could potentially result in some increases in *Vibrio* concentrations, they did not appear to affect the overall pattern of results. This was tested with logarithmic regression analysis that showed no relationship with arrival temperature or delivery time (R^2^ = 0.001 and 0.003 for *V. vulnificus* and *V. parahaemolyticus*, respectively) and in the worst-case 15 February 2017 sample there were no *V. vulnificus* detected and *V. parahaemolyticus* were typical for that time of the year (919 MPN g^−1^). Checks were also made that the *V. vulnificus* peak concentrations that the study focused on did not come from samples with high arrival temperatures or delivery times; however, no anomalies were observed, so results from all samples were included in the study.

### Laboratory testing

*Vibrio vulnificus* and *V. parahaemolyticus* concentrations in the oysters were determined as described previously (21). Briefly, samples of 12 live oysters were shucked, homogenised in their own intervalvular fluids and diluted in Alkaline Peptone Water in a three-tube MPN format and incubated at 35°C for 24 h. The presence of the *Vibrio* species was determined in each well using quantitative PCR (qPCR), and MPNs calculated. Salinity of the seawater samples from each site was determined using a refractometer (Atago S/Mill, Japan).

### Data analyses

Genstat statistical software (22) was used to carry out Analysis of Variance (ANOVA) to compare log transformed *Vibrio* concentrations in oysters from different sites. Where results showed normal distribution, least significant differences (LSD, P = 0.05) were used to compare results. Attempts were made to establish relationships between detected *V. vulnificus* concentrations in oysters and total, minimum and/or mean values of rainfall, seawater temperature and salinity and over the 24, 48, 96, 144, 168, 240, 288, 312, 336 h before the sample was collected, plus with lag times (e.g. salinity between 144 and 216 h (6–9 days) before sample collection). Random Forests and then Best Subsets Regression by year or across all years were tried to give an indication of the most important variables. A structural equation model (fitted in R using the lavaan package) was then applied to all of the important variables. All variables were fitted to the whole model initially, then those terms that were not significant were gradually dropped (‘backward elimination’) to achieve a best fit model. Additionally, rates of decline after peaks in concentration were determined in an attempt to consider how long growers might need to wait after a *Vibrio* event before concentrations returned to normal.

## Results

### Harbour A

Only one oyster sample was collected each fortnight in 2016 and the results from Harbour A are presented in Figure 3. ANOVA of the *V. vulnificus* results from the different Harbour A sampling sites across all the 2017–19 sampling dates, blocking by harvest date, did not show significant differences (F = 0.554) between sites, so averages of the three sites with the calculated LSD of 0.752 log_10_ MPN g^−1^ were used to compare the log concentrations of *V. vulnificus* at different times (Figures 4–7). In contrast, for *V. parahaemolyticus*, results suggested that, on average, Site 7 might have had marginally lower *V. parahaemolyticus* counts than Sites 11 and 12 (logarithmic means across all samplings = 373, 575 and 456 MPN g^−1^, respectively, log_10_ LSD = 0.170). However, although the LSD (0.170 log_10_ MPN g^−1^) suggested a difference, the ANOVA F statistic did not confirm significant differences (F = 0.094). As *V. parahaemolyticus* results showed similar patterns to those published previously with concentrations being dominated by water temperatures (20, 19) these results are not further detailed here but are available in Supplementary File. In contrast, we detected many more Pacific oyster samples containing *V. vulnificus* than the 2 of 13 samples (maximum concentration = 0.36 g^−1^) that had previously been published for Harbour A and higher concentrations than had been observed in any other harbour (13; see Figure 1). Therefore, here we are examining the *V. vulnificus* results from this harbour in depth to better understand their relationship to environmental parameters (rainfall, seawater salinity and temperature). Seawater salinity data had a lot of noise, and temperature in the shallow estuarine waters often fluctuated markedly over a 24-h period so, to better visualise the data, mean results from the previous 24 h were plotted for each 15-min interval recording in Figures 3 to 6.

**Figure 3.**
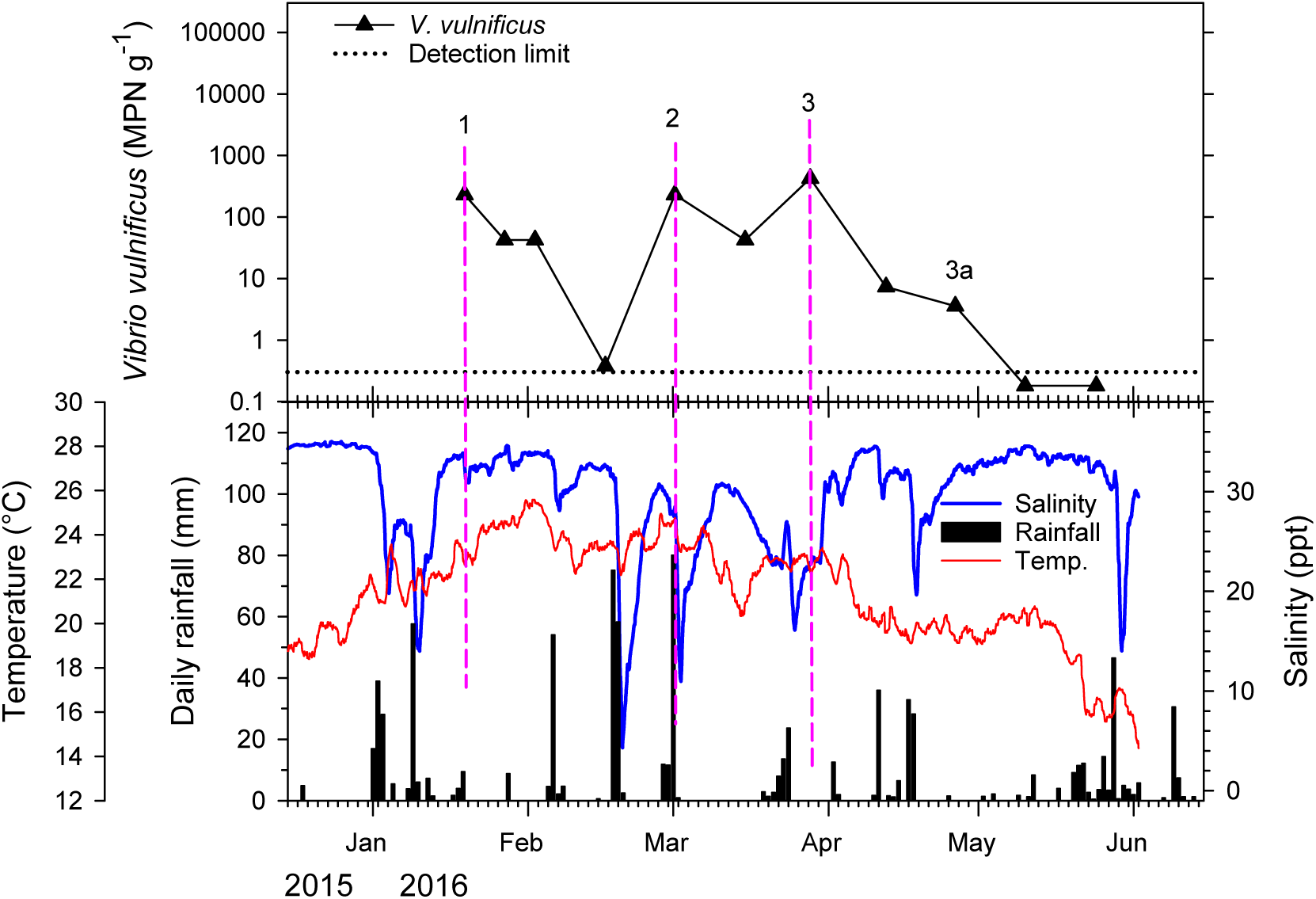
2016 concentrations of *Vibrio vulnificu*s in Pacific oysters from Harbour A during the summer months as related to parameters of the oyster growing environment (rainfall, mean seawater salinity and temperature over the previous 24 h). Numbered vertical dashed lines identify peaks in *V. vulnificus* numbers. A single sample of 12 oysters was tested on each occasion. Plotted salinities and temperatures are the means for the previous 24 h.

**Figure 4.**
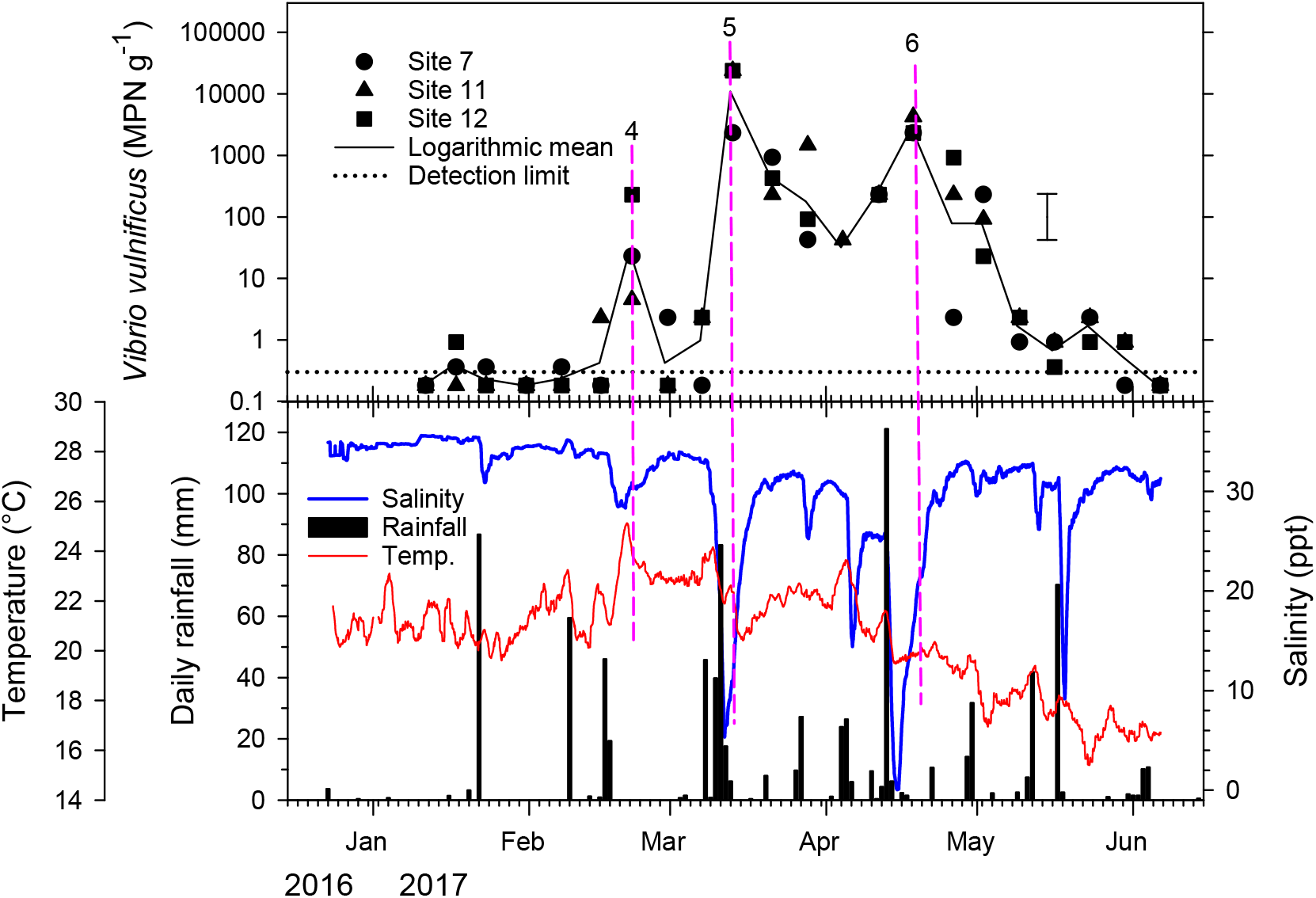
2017 concentrations of *Vibrio vulnificus* in Pacific oysters from Harbour A during the summer months as related to parameters of the oyster growing environment (rainfall, mean seawater salinity and temperature over the previous 24 h). A single sample of 12 oysters was tested from each sampling site (Figure 2) on each occasion, and as differences between each site were not significant, changes in mean concentrations of the three neighbouring sites (7, 11, and 12) are plotted. Error bar represents least significant difference (LSD) of the logarithmic means (P = 0.05) as calculated over the summers of 2017–19 (Figures 4–6). Numbered vertical dashed lines identify spikes in *V. vulnificus* numbers. Plotted rainfall, salinities and temperatures are means over the previous 24 h.

Over the course of the 3-year study (2017–19) with weekly sampling of the three sites over the summer months, and with fortnightly sampling in 2016 at a single site, nine occasions were observed in which *V. vulnificus* concentrations spiked by more than the LSD between sampling occasions (Figures 3–7). The first sampling in 2016 (Peak 1 in Figure 3) probably represented another such peak and there was another lesser peak (Peak 10 in Figure 5) that suggested a similar response to environmental changes as will be describe below. In 10 of the 11 peaks, the increases in *V. vulnificus* numbers followed a period of sustained and/or heavy rainfall resulting in seawater salinities dropping to below 20‰ compared with the average of 32‰ for the harbour.

**Figure 5.**
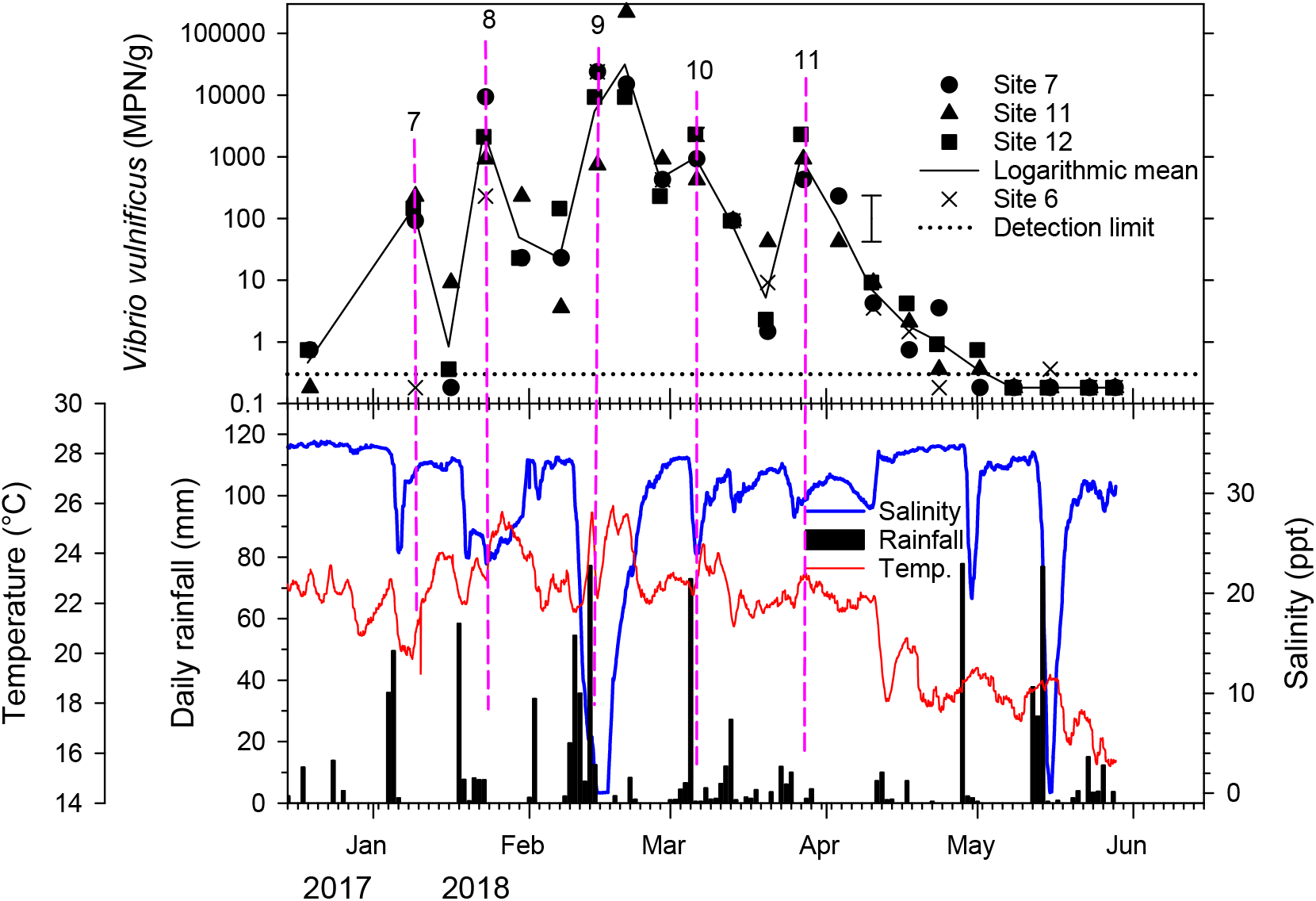
2018 concentrations of *Vibrio vulnificus* in Pacific oysters from Harbour A during the summer months as related to parameters of the oyster growing environment (rainfall, mean seawater salinity and temperature over the previous 24 h). A single sample of 12 oysters was tested from each sampling site (Figure 2) on each occasion, and as differences between each site were not significant, changes in mean concentrations of the three neighbouring sites (7, 11, and 12) are plotted. Error bar represents least significant difference (LSD) of the logarithmic means (P = 0.05) as calculated over the summers of 2017–19 (Figures 4–6). Numbered vertical dashed lines identify spikes in *V. vulnificus* numbers. Plotted rainfall, salinities and temperatures are means over the previous 24 h.

In 2016, the first sample collected on 19 January showed an elevated (>5 MPN g^−1^) *V. vulnificus* concentration of 231 MPN g^−1^ (Figure 3, Peak 1). This had been after a prolonged period of rainfall in early January (83 mm between 1 and 3 January) and very high rainfall on 9 January that had resulted in the 24-h average salinity dropping to 14.2‰. Salinities had returned to normal by the time of sampling; however, *V. vulnificus* concentrations remained elevated through to 2 February (42 MPN g^−1^) but had dropped to the detection limit (0.36 MPN g^−1^) by 16 February. The next observed *V. vulnificus* peak (Figure 3, Peak 2) on 1 March also occurred after a period of heavy rainfall on 18 and 19 February (total of 133 mm) resulting in the average salinity dropping to just 4.3‰ and another rainfall event on 29 February (80 mm, 11.0‰). By 15 March, the average daily salinity was above 28‰ and *V. vulnificus* had dropped back to 42 MPN g^−1^. Then there was another smaller rain event (52 mm between 19 and 24 March) bringing the average daily salinity down to 16.1‰ by 7:00 am on 25 March, and the 29 March sample had a significantly increased *V. vulnificus* concentration of 424 MPN g^−1^ (Figure 3 Peak 3). A fortnight later on 12 April, the salinity had risen back above 32‰ and *V. vulnificus* concentration had dropped to 7.3 MPN g^−1^. There was then another rain event resulting in salinity dropping to 20.0‰ on 18 April. No increase in concentration was observed but the downward trajectory might have been slowed at point 3a on Figure 3. The lack of an increase might be related to lower water temperatures that had dropped to a daily mean of 19.6°C compared with at least 22.8°C when the other peaks occurred.

In 2017, 87 mm of rain fell on 22 January but seawater temperatures were still below 20°C and the average salinity only dropped from 35.1 to a minimum of 32.0‰ at 6:15 pm on 22 January and no peak in *V. vulnificus* was detected. However, there was further significant rainfall on 9 February (59.5 mm) and again on 16 and 17 February (66 mm over the 2 days), and this time average salinity dropped to 28.5‰ on 19 February when seawater temperature was 24.9°C. A small peak in *V. vulnificus* concentration (average 23 MPN g^−1^) was then observed when sampled on 21 February. The next major rainfall event occurred on 8–13 March when a total of 193 mm of rain fell, flooding the harbour and causing the daily salinity to drop as low as 5.3‰ on 11 March. This resulted in a massive peak of *V. vulnificus* concentrations, reaching a mean of 10,900 MPN g^−1^ when sampled on 13 March (Figure 4, Peak 5). Concentrations then declined over the next 3 weeks (42 MPN g^−1^ on 4 April) despite some rainfall on 26–27 March (37 mm) and 2–6 April (57 mm). However, following the latter, the average salinity dropped to 14.3‰ on 6 April resulting in concentrations of 230 MPN g^−1^ on 13 April. Then there was very heavy rain (121 mm) on 13 April with average salinity reaching a minimum of 0.06‰ on 15 April and, when next sampled on 18 April, *V. vulnificus* concentrations had risen to a mean of 2840 MPN g^−1^ (Figure 4, Peak 6). There was further heavy rainfall in May (70 mm on 17 May) resulting in average salinities of 9.1‰ on 18 May. However, the maximum *V. vulnificus* concentration detected when oysters were next sampled on 23 May was only 2.3 MPN g^−1^, probably reflecting the lower temperature around this time (c. 17°C).

In 2018, five peaks in *V. vulnificus* were observed, all following rainfall and reduced salinities with seawater temperatures of ≥ 20°C (Figure 5). As well as the usual sampling sites (7, 11 and 12 in Figure 2), 14 samples were collected from Site 6, which was further out in the harbour and might be less subject to reduced salinity. The salinities recorded from Site 6 seawater samples collected on the day of sampling were only slightly higher than those from the other three sites on the same days (mean of 28.9 compared with a mean of 26.9) and low salinities were also recorded at Site 6 (e.g. 5‰ at Site 6 compared with 3, 3 and 2‰ at Sites 7, 11 and 12, respectively, on 14 February. *V. vulnificus* concentrations in oysters from Site 6 were usually similar to those of the other sites (Figure 5), e.g. they were all at 92 MPN g^−1^ on 13 March. The exception was on 9 January when the concentration at Site 6 was only 0.18 MPN g^−1^ compared with the logarithmic average of 146 MPN g^−1^ from the other three sites, although the salinity on the day was 35‰ at both Sites 6 and 7. To avoid any effects of the Site 6 results, the logarithmic averages plotted in Figure 5 are of the same sites available in Figures 3, 4 and 5, excluding Site 6. Peak 7 on 9 January occurred after 87 mm rain on 4–6 January, although the average salinity only dropped to 24.1‰ on 6 January. In Peak 9, the highest concentration of *V. vulnificus* (221,000 MPN g^−1^) that has ever been detected in New Zealand shellfish was recorded. For Peak 10, 73 mm rain fell on 5 March and salinities had dropped to 26‰ by the time the samples were collected on 6 March. This halted the downward trend in *V. vulnificus* after Peak 9 with a slight increase from 478 MPN g^−1^ on 27 February to 966 MPN g^−1^. There were rainfall events resulting in reduced salinity in late April and mid-May but because seawater temperatures were below 14°C these events did not cause *V. vulnificus* concentrations to rise. As well as reduced salinity, rainfall may have resulted in an influx of nutrients into the harbour. Nutrient levels were not monitored.

There was a much drier summer in 2019 and the only times salinity was recorded to drop below 29‰ were on 2–4 December, 25–26 December and 9–15 April dropping to 28.2, 26.7 and 25.2‰, respectively (Figure 6). However, mean temperatures were all below 20°C at those times and no *V. vulnificus* peaks were detected.

**Figure 6.**
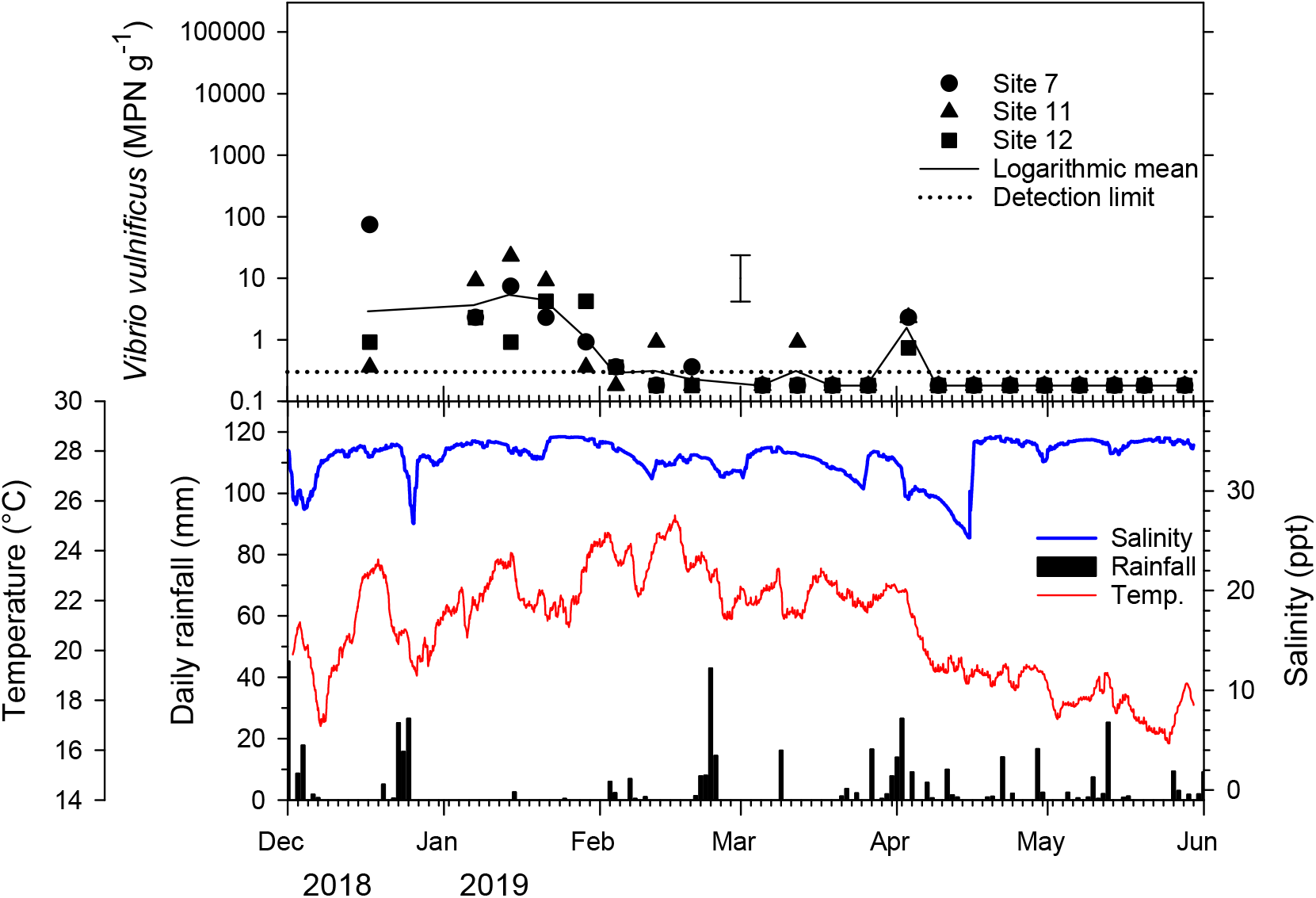
2019 concentrations of *Vibrio vulnificus* in Pacific oysters from Harbour A during the summer months as related to parameters of the oyster growing environment (rainfall, mean seawater salinity and temperature over the previous 24 h). A single sample of 12 oysters was tested from each sampling site (Figure 2) on each occasion, and as differences between each site were not significant, changes in mean concentrations of the three neighbouring sites (7, 11, and 12) are plotted. Error bar represents least significant difference (LSD) of the logarithmic means (P = 0.05) as calculated over the summers of 2017–19 (Figures 4–6). Numbered vertical dashed lines identify Peaks in *V. vulnificus* numbers. Plotted rainfall, salinities and temperatures are means over the previous 24 h.

**Figure 7.**
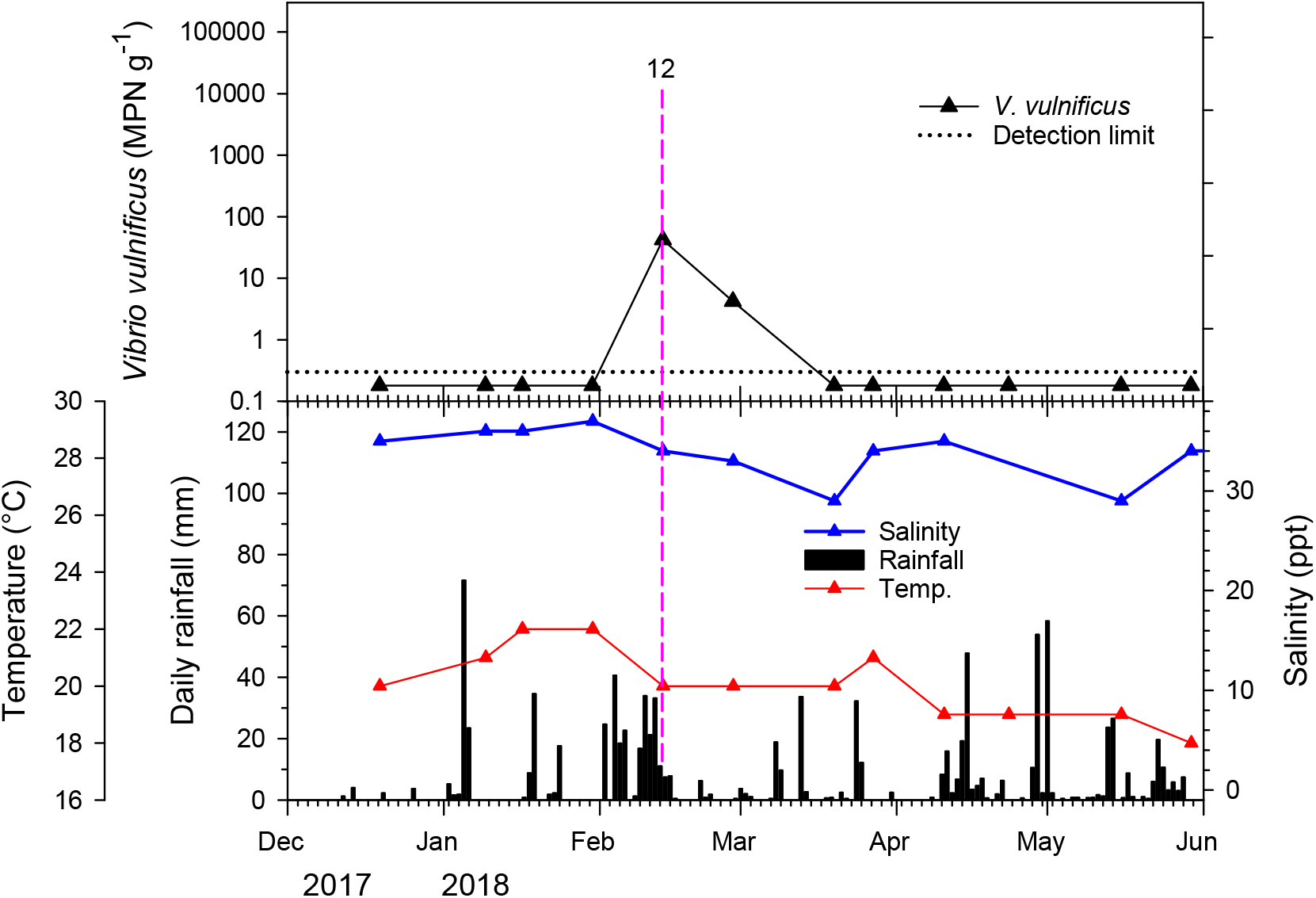
2018 concentrations of *Vibrio vulnificus* in Pacific oysters from Harbour D during the summer months as related to parameters of the oyster growing environment (rainfall, growing water salinity and temperature). Numbered vertical dashed line identifies a peak in *V. vulnificus* numbers.

### Harbours C and D

Oysters from Harbours C and D did not show the same peaks in *V. vulnificus* concentrations as were found in Harbour A. The maximum number from Harbour C was only 2.3 MPN g^−1^ and there was only one occasion when this was exceeded in Harbour D, reaching 42.4 MPN g^−1^ (Figure 8, peak 12).

**Figure 8.**
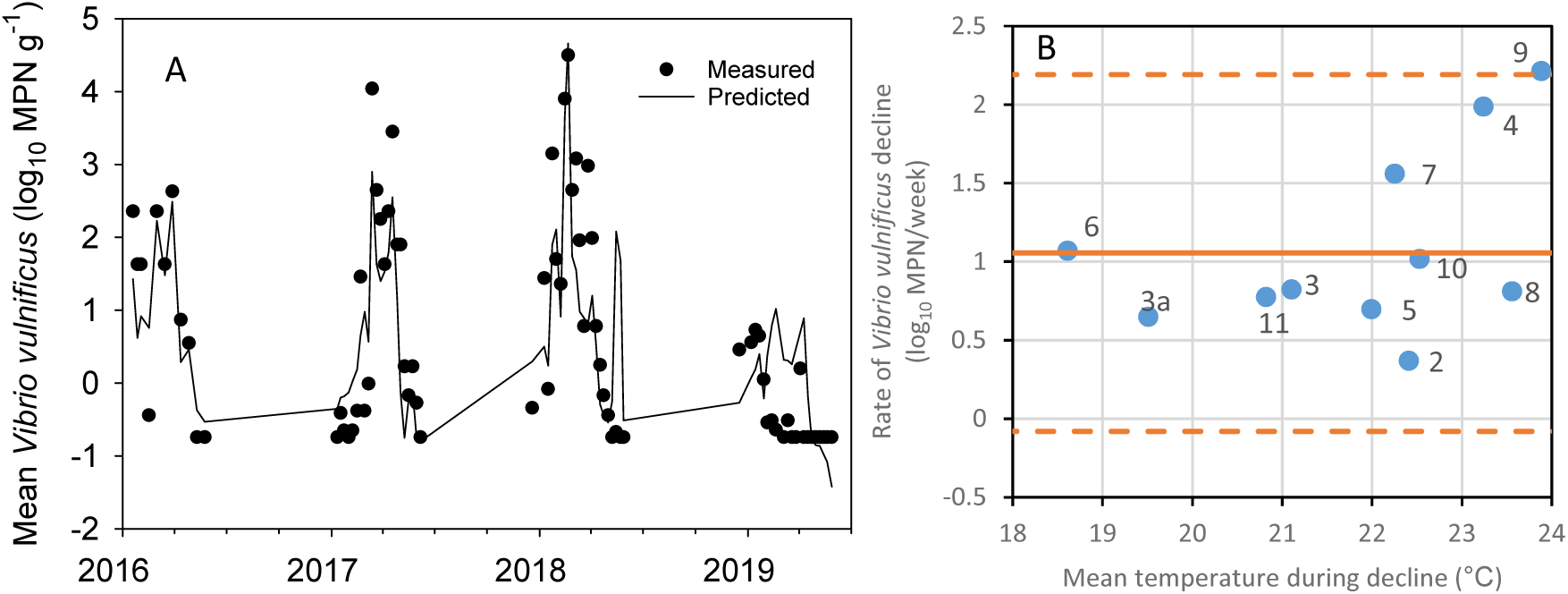
Modelling *Vibrio vulnificus* concentrations in Pacific oysters in Harbour A over 4 years. A. Measured versus predicted concentrations from Equation 2 (Log_10_ *V. vulnificus* per g of oysters = 0.883 + 0.267 x mean temperature 144 to 168 h ago – 0.095 x mean salinity 0 to 24 h ago – 0.098 x mean salinity 168 to 192 h ago). B. Rates of decline in concentration at different seawater temperatures. Labels of the data points are the peak numbers identified in Figures 3–6. Solid horizontal line is the mean rate of reduction with the dashed lines representing the 95% confidence intervals.

Although only collected on a fortnightly basis, the spot temperature and salinity results collected at Harbour D (mean = 19.9°C and 34.6 ‰) and Harbour C (means = 20.9°C and 33.7‰) were not much different from those recorded from the weekly Harbour A water samples (means = 21.1°C and 31.5‰). However, there was no evidence of big drops in salinity after rainfall with minimums of 29 and 25‰, respectively, in Harbour D and Harbour C compared with 2‰ in Harbour A. Peak 12 detected on 13 February 2018 in Harbour D (Figure 8) occurred after a period of prolonged rainfall (224 mm since 2 February) that may have resulted in reduced salinity. However, on the day of sampling when a further 11 mm of rain fell, the salinity was only 34‰ compared with the 37‰ recorded from the previous sampling period and 33‰ from the following one.

### Predictive modelling

When trying to establish relationships between environmental parameters and *V. vulnificus* concentration, random forests did not give a clear indication of which variables were important. Best subsets regression suggested two- or three-variable models were best for individual years and the best predictions used: mean temperature of the previous 24 h, precipitation (previous 168 h) and perhaps minimum temperature (previous 24 h) for 2016; minimum salinity (previous 24 h), and either minimum salinity (previous 96 h) or precipitation (previous 120 h) for 2017; mean temperature (previous 24 h) and minimum salinity (previous 96 h) for 2018; and mean temperature (previous 24 h) and minimum salinity (previous 24h) for 2019. Including lag times in the best subsets regressions, salinity in the previous 24 or 48 h and for the salinity between 12 and 13 days (288–312 h) before sampling were best for 2017 and 2019, whereas temperature between 6 and 9 days plus precipitation in the previous week (168 h) was best for 2018. When looking across all 4 years, best subsets regression suggested that four- or five-variable models were best, accounting for 65–67% of the variance with the best variables being: minimum salinity (previous 288–312 h); minimum salinity in either the 24 or 24–48 h previous; minimum salinity in either the 72–96 or 96–120 h before sampling and minimum salinity in the previous 96 h.

Applying the structural equation model initially gave Equation 1 that accounted for 65% of the variance.

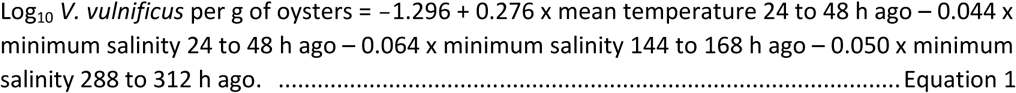

However, we were concerned that minimum salinity might not be the most reliable measure to consider since one aberrant reading could have undue influence on the prediction (e.g. if the sensor was exposed to air and rain during low tide in an intertidal setting). The exercise was therefore repeated using mean salinity rather than minimum. Using best subsets regression, the best fitting multi-variable models had four variables and accounted for about 60–62% of variation, with mean salinity in the last 24 h and mean temperature 144–168 h ago in all the best models along with mean salinity either 144–168 h ago or 168–192 h ago, and minimum salinity either 48–72 h ago, 72–96 h ago, or 96–120 h ago.

In fact, the best of the three-variable models account for about 59–60% of variation, using mean salinity in the last 24 h and mean temperature 144–168 h ago and mean salinity either 144–168 h ago or 168–192 h ago (the latter was slightly better with 60% variance accounted for, rather than 59%). The three-variable model is:

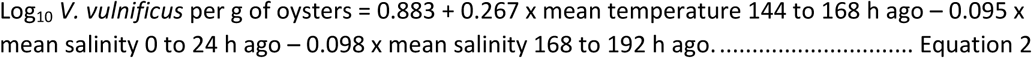

As a cross check, we plotted the predicted values from this equation against the actual Harbor A data over the 2016–19 years (Figure 8). The model got the overall shape correct for 2016–18. It also predicted 2019 to have lower concentrations, although the shape was not quite correct when these low numbers were present.

As well as attempting to model concentrations based on environmental parameters, we paid particular attention to rates of decline in concentration after a concentration peak. Rates of decline were determined for each peak in Figures 3–6 and plotted against the mean seawater temperature during the period of the decline (Figure 8B). Rates of decline averaged 1.05 log_10_ MPN week^−1^ but ranged from 2.214 to 0.368 log_10_ MPN week^−1^. All three of the highest rates of decline (Peaks 7, 4 and 9, all with rates of over 1.5 log_10_ MPN week^−1^) occurred at seawater temperatures above 22°C, but slower rates also occurred above 22°C including the lowest rate of decline (0.368 after Peak 2) that occurred when mean water temperature was 22.4°C. Examining the rates of decline in relation to salinity and date/year did not show any particular trends.

## Discussion

Harbour A (Figure 2) contains a relatively large mass of estuarine water with only a small entrance to the harbour to facilitate mixing with the marine waters of the Pacific Ocean. The tidal exchange between the two bodies of water is relatively small, with New Zealand Marine Charts (https://wetmaps.co.nz/) recording tidal flows of just 1.7 knots at the narrow entrance to the harbour with a depth of just 15 m at the shallowest part. Despite this, unlike many other estuarine systems in the world (e.g. Chesapeake Bay, USA), even in the upper reaches of the harbour where this study was carried out, salinities are typically above 32‰, well above optimum salinities for the growth of *V. vulnificus* (23). Under these conditions, low numbers of *V. vulnificus* could still be detected in oysters in Harbour A while seawater temperatures were above 20°C. For example, concentrations averaged 4.5 MPN g^−1^ between 7 and 21 January 2019 when there had not been any rainfall (Figure 6). However, as detailed in the results, this study shows that after periods of heavy rainfall the salinities at intertidal oyster farms in the shallow upper reaches of the harbour dropped dramatically, at times reaching 0.0‰ (freshwater). At such times, when seawater temperatures were above 18°C, *V. vulnificus* concentrations increased dramatically. By sampling Site 6 (Figure 2) we considered whether this effect might be lesser further out in the harbour, closer to the ocean.

However, salinities at this point were only about 2‰ higher than the other sites and, with the exception of Peak 7 (Figure 5), *V. vulnificus* concentrations were similar.

There was a clear relationship between high rainfall that resulted in reduced salinity and high concentrations of *V. vulnificus* in oysters. Ten of the 11 peaks identified in Figures 3–6 could be attributed to low salinity events that occurred when seawater temperatures were above 16°C. In contrast, in 2019 when rainfall was low and salinities were relatively constant, we did not detect any peaks in *V. vulnificus* concentration. The one peak observed in Harbour D also occurred after significant rainfall. Low salinity may have gone undetected in that harbour with only fortnightly monitoring. Most of the peaks that occurred after a significant rainfall event caused 24-h salinities to fall below 25‰; however, Peak 4 (which was small) and Peak 11 occurred after events giving salinities closer to 30‰. Other causative mechanisms may have been at play in these cases and in the small peak observed in Harbour D. Peaks in *V. vulnificus* concentrations usually only occurred when seawater temperatures were above 20°C, but Peak 6 occurred when temperatures had been about 19°C. This lower temperature at Peak 6 might explain why this peak was lower than Peaks 5 and 9 despite the low salinity. In cases when low salinity events occurred at seawater temperatures below 19°C, no rise in *V. vulnificus* concentrations was observed. In three cases, low salinity events at temperatures between 19 and 20°C also did not cause rises (in April 2016, and December and April 2019).

Overall, we can say that rainfall resulting in salinities of less than 24‰ at seawater temperatures of over 20°C always caused increased *V. vulnificus* concentrations in oysters, while salinities of less than 30‰ combined with seawater temperatures above 19°C also often caused such increases.

Nigro (24) also recorded increased *V. vulnificus* abundance in seawater after a major rain event had resulted in salinities dropping from normal (above 30‰) to 13‰ and remaining low for 16 h. She suggested that “sustained depressed salinities may present an increased risk for *V. vulnificus* infection.” Bullington *et al*. (25) also observed that *V. vulnificus* growth increased after summer rain storms. However, they demonstrated that as well as salinity and temperature effects, increased nutrients measured as fluorescent dissolved organic matter and inorganic nutrients, particularly silicates, also contributed to increased *V. vulnificus* density in the seawater. Nutrients were not measured in the current study so effects of reduced salinities from the influx of fresh rainwater cannot be distinguished from any increase in nutrients washed into the harbour from rainwater running off from the land. However, as *V. vulnificus* grows poorly at salinities above 30‰, the reduced salinities must have had an influence on the peaks in *V. Vulnificus* numbers.

Although low salinity and warm seawater clearly play a role in increasing *V. vulnificus* in oysters, the relationship is by no means straightforward, as shown by the complexity of Equations 1 and 2 and the fact that they only accounted for 60% of the variance in concentration. Other factors might be involved, including the amount of nutrients in the seawater. It is not known why the relatively high concentrations of *V. vulnificus* occasionally found in Pacific oysters have not caused food-borne illnesses. It may be that people with underlying health issues have taken heed of public health warnings to not eat raw shellfish or it may be related to the different strains present in New Zealand compared to parts of the world that have regular outbreaks (Cruz *et al*. 2016). However, should *V. vulnificus* ever become of public health concern in New Zealand, using data from temperature and salinity monitoring buoys with Equation 2 would give a pre-emptive warning of when *V. vulnificus* concentrations are likely to increase. Based on the data generated in this study, salinities above 30‰ or temperatures below 19° are unlikely to cause peaks in *V. vulnificus* concentrations.

However, concentrations seem certain to increase when salinities are less than 24‰ with temperatures of over 20°C, and they may also increase at salinities of up to 30‰ and temperatures down to 19°C. With climate change predicted to increase the frequency and severity of flooding in various parts of New Zealand, it is likely that high concentrations of *V. vulnificus* will occur more frequently in other harbours.

If *V. vulnificus* was to be regulated for public health reasons, an important practical question would be, once concentrations increase to any nominally unacceptable level, how long would it take for them to return to normal low levels? At 1.05 log_10_ week^−1^, the average rate of decline was relatively slow compared with the likes of faecal coliform depuration in Pacific oysters where at 20°C, 2 log_10_ reductions were achieved within 24 h (26). Further, because the rate of decline was highly variable, microbiological testing would also need to be carried out to confirm how long a peak in *V. vulnificus* concentration might take to return to any predetermined acceptable concentration.

The current study provides a greater understanding of the population dynamics of *V. vulnificus* in New Zealand Pacific oysters, which will help with risk assessment and management should the strains of the organism present in New Zealand be found to cause foodborne illness.

## Acknowledgements

We are grateful to the members of the New Zealand oyster industry for the collection of samples and access to buoy data. Without their help the study could not have been conducted.

We also wish to thank Michael Quach and Nicola Wei for technical assistance.

The project was funded by the New Zealand Ministry of Business, Innovation and Employment (MBIE) through the Cawthron Institute’s Seafood Safety programme, contract CAWX1801.

*V. parahaemolyticus* 2016 numbers presented in Figure 3 were collected in a monitoring programme funded by the New Zealand Ministry for Primary Industries (MPI), contract 33253.

## Conflict of interest

No conflict of interest declared.

